# Liquid-state surface fermentation as a platform for pure mycelium materials production

**DOI:** 10.64898/2026.10.26.761940

**Authors:** Jara Saluena Martin, Piet Dimitriadis, Soufian Attalibi, Annah-Ololade Sangosanya, Rafael Philippini, Razieh Shafiei-Alavijeh, Joost Brancart, Elise Elsacker, Eveline Peeters

## Abstract

Liquid-state surface fermentation (LSSF) is emerging as an alternative to submerged and solid-state fermentation for producing pure mycelium materials (PMM), yet a systematic framework linking culture geometry, substrate consumption, inoculum preparation and material properties is lacking. Here, we characterised LSSF with *Trametes versicolor* across culture volumes and inoculation strategies. The biomass accumulation model described a trade-off between colonisation speed and biomass accumulation. Smaller volumes, with a higher specific interfacial area, colonised the surface earlier and converted glucose more efficiently (up to 0.64 g g⁻¹), but depleted it prematurely and produced less biomass. After pellicle formation, glucose was consumed at a common volumetric rate (∼1.0 g L⁻¹ d⁻¹), having glucose availability as a limiting factor for pellicle densification. In deeper cultures, much of the glucose consumed early was not recovered in the pellicle, indicating competition from submerged growth. Backlit transmittance imaging provided a non-destructive estimate of pellicle biomass and of its spatial heterogeneity. Inoculum source and processing determined material performance: liquid-derived inocula and a shaken incubation step increased biomass, an additional blending step produced more homogeneous and stronger pellicles. These results provide process-level design rules for LSSF and a basis for its application to PMM production.

**Graphical Abstract:** 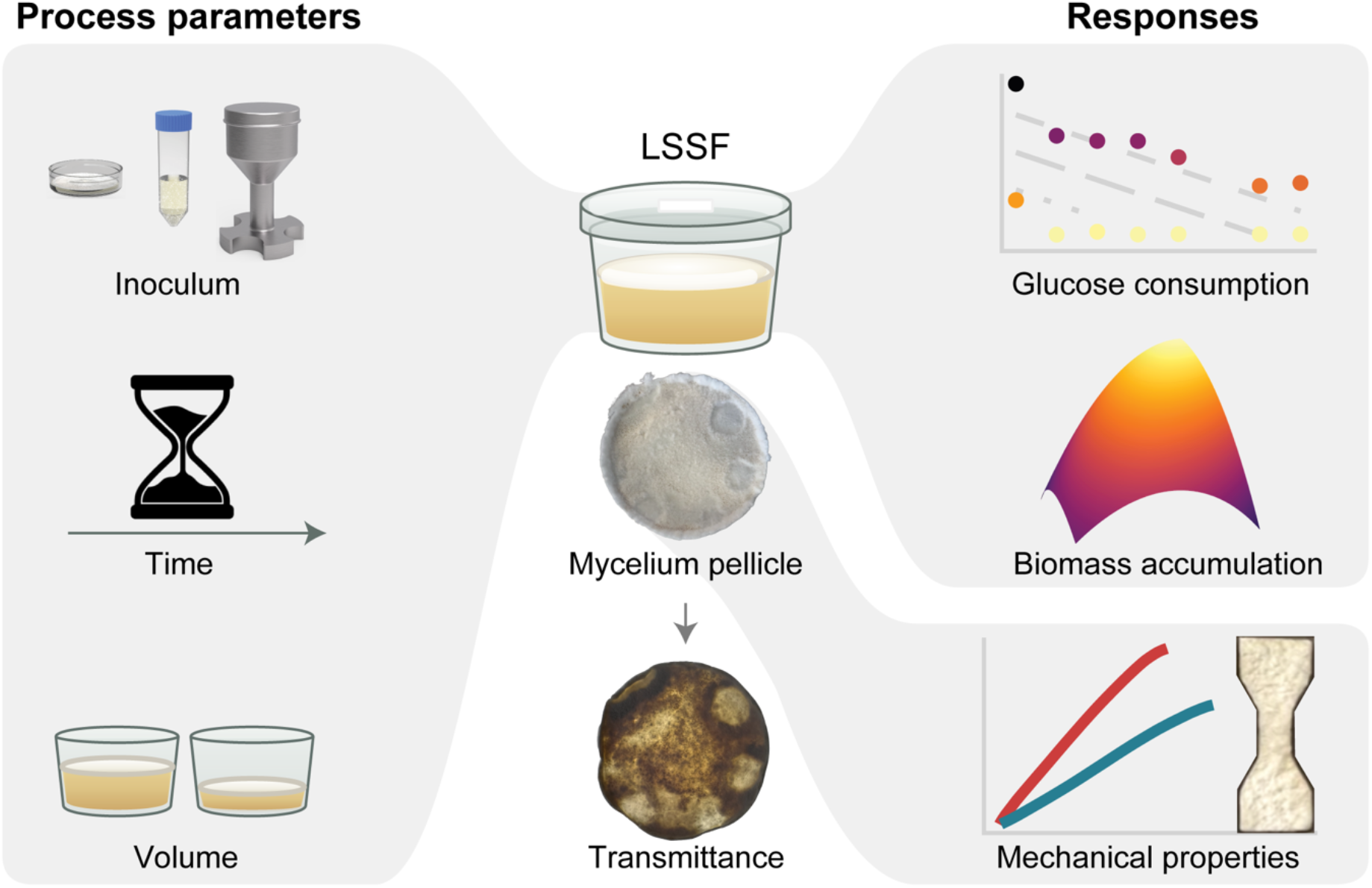

## Introduction

Mycelium, the vegetative structure of filamentous fungi, can be cultivated to produce materials that offer sustainable alternatives to fossil-based plastics and animal-derived materials. Mycelium materials can be produced from low-value agricultural or industrial side streams, offering opportunities for lower-carbon material production (Livne et al., 2022; Alaux et al., 2024). White-rot basidiomycete fungi are commonly used for these materials because they efficiently colonise lignocellulosic substrates through dense filamentous growth. In contrast to mycelium-based composites (MBCs), in which hyphae bind substrate particles into a cohesive matrix (Grimm and Wösten, 2018; Elsacker et al., 2019), pure mycelium materials (PMMs) consist primarily of fungal biomass, obtained by separating the mycelium from the nutrient source (Gandia et al., 2021; Vandelook et al., 2021).

PMMs can be produced through three main cultivation approaches: liquid-state submerged fermentation (SmF), liquid-state surface fermentation (LSSF) and solid-state surface fermentation (SSSF). In SmF, fungal biomass develops submerged in a liquid nutrient medium, often as pellets or dispersed mycelium (d’Errico *et al*., 2024; Wijayarathna *et al*., 2026). In SSSF, mycelium grows predominantly over a solid substrate surface, allowing the aerial biomass to be harvested. In LSSF, mycelium develops as a cohesive pellicle at the air-liquid interface, combining a liquid nutrient phase with static surface growth (Elsacker *et al*., 2023). LSSF enables fungal cultivation without mechanical agitation or homogenisation, which may reduce energy consumption and equipment fouling. In addition, the liquid phase offers flexibility in feedstock selection, including both soluble organic substrates and submerged lignocellulosic solids.

LSSF has been used in other biotechnological processes, including kombucha production by mixed yeast-acetic acid bacterial consortia and citric acid production by the ascomycete *Aspergillus niger*, although these processes target fermented liquids or secreted metabolites rather than the fungal pellicle itself (Tran *et al*., 2022; Singh Dhillon *et al*., 2011). Because these systems rely on aerobic microbial activity, oxygen availability is a key determinant of process performance. Under static conditions, dissolved oxygen is determined by the surface area exposed to air (Hornung et al., 2006; Rodrigues et al., 2026). Thus, larger surface areas typically enhance oxygen transfer across the culture volume and promote microbial growth (Tran *et al*., 2020). However, in the context of LSSF-based mycelium material production, the interplay between surface area and culture volume remains largely unknown.

Beyond culture geometry, the production of mycelium materials can be tailored by several other process variables to improve yield and achieve specific material properties, including texture, thickness and colour (Appels *et al*., 2019; Antinori *et al*., 2020). Strain selection, cultivation setup, bioprocess conditions and post-processing steps have been shown to be key design choices (Haneef *et al*., 2017; Alemu *et al*., 2022). For example, the state of the fungal inoculum has been proved to have a major impact on fermentation performance in fungal SmF as well as SSSF for MBC (Liu *et al*., 2016; Saluena Martin *et al*., 2026). Liquid inoculum preparation typically involves biomass disruption methods, such as blending or sonication, and may include sequential incubation steps that allow biomass regeneration before inoculation of the production culture. Although such protocols can affect inoculum dispersion and growth performance, their impact on biomass yield and material properties of PMM remains poorly understood. Moreover, the role of inoculum source and preparation in LSSF-based PMM production has not yet been systematically investigated.

In this study, we used the polypore basidiomycete *Trametes versicolor* to establish LSSF as a reproducible platform for producing pure mycelium mats. We set out to analyse biomass formation and substrate consumption across different culture volumes and incubation times and to compare inoculum preparation strategies differing in biomass source and mechanical disruption approaches on the mechanical properties of the material. As such, we aim to move beyond feasibility demonstrations of LSSF and to provide process-level design rules for PMM production.

## Experimental procedures

### Strain and pre-culture conditions

*Trametes versicolor* strain VUB-SV1 was maintained on solid potato dextrose agar (PDA; Condalab, Spain) Petri plates at 4 °C.

Fungal cultivation was performed in Complete Yeast Medium (CYM) containing 2% D-glucose monohydrate, 0.3% peptone, 0.2% yeast extract, 0.05% MgSO_4_, and 9 mM phosphate buffer (pH ∼ 6.8). For solid pre-cultures, 1.5% agar was added to CYM in 60-mm Petri plates (Greiner, Austria). For liquid pre-cultures, CYM was supplemented with 60 μg/mL kanamycin and dispensed into either 50-mL Falcon tubes or 250-mL Erlenmeyer flasks. Pre-cultures were inoculated with 5-mm mycelium plugs excised from the periphery of freshly grown colonies or with homogenate from a previous pre-culture. Plates and Falcon-tube pre-cultures were incubated in static and dark conditions at 25 °C for five days, unless otherwise stated. Pre-cultures in Erlenmeyer flasks were incubated in dark conditions at 28 °C while shaking at 200 rpm (New Brunswick Innova 40/40R, Eppendorf, Germany) during 24 hours.

### Liquid-state surface fermentation

Homogenisation of mycelial biomass – derived from either solid or liquid pre-cultures – was performed in 100 mL CYM with a 360 mL stainless steel semi-micro blending cup (Eberbach, USA) and a blending base (Waring, USA). Each homogenisation step consisted of two 5-second blending cycles at speed 5, separated by a 30-second rest period. Homogenates from equivalent treatments were pooled to decrease variability in the production culture.

LSSF was performed in 90 x 60 mm round polypropylene microboxes sealed by filtered lids (O95/60+OD95, SacO2, Belgium). By default, microboxes were filled with 90 mL CYM and 18 mL inoculum. For experiments using total culture volumes different from 108 mL, the inoculum-to-medium volume ratio was kept constant at 1:5 (v/v). Microboxes were incubated statically in the dark at 28 °C for 7 to 14 days.

Mycelial mats that developed on the surface of the medium were removed from the microboxes, gently rinsed with distilled water to remove the bottom exopolysaccharide layer, lightly blotted with paper towels, and weighed to determine fresh biomass. Materials were then glycerol-treated by soaking each specimen in 50 mL of a 10% glycerol (Sigma-Aldrich, USA) solution for 3 to 4 hours and air-dried.

To establish the relationship between wet and dry biomass, mycelium pieces of varying sizes were excised from freshly harvested mats, weighed, lyophilised using a 2.5-L freeze dryer (Labconco, USA), and weighed again to determine dry mass.

### Chemical analysis of spent medium

At harvest, spent medium pH was measured using a calibrated digital pH meter (C831, Consort, Belgium). For quantification of reducing sugars, aliquots were centrifuged at 12000 *g* during 1 minute twice to remove exopolysaccharides, followed by a 3,5-dinitrosalicylic acid (DNS) colorimetric assay (Miller, 1959). This assay was performed in a 96-well microplate, as described by Garcia-Gonzalez et al. (2019) with slight modifications. The DNS solution contained 30% potassium sodium tartrate, 1.6% NaOH, and 1% DNS reagent. After centrifugation, supernatants were diluted 1- to 10-fold depending on initial screenings and subsequently 2-fold with DNS solution in a total volume of 100 μL. Microplates were covered with Microseal ‘B’ films (MSB1001, Bio-Rad Laboratories, USA) and incubated at 90 °C for 20 minutes. After an additional 2-fold dilution in water, absorbance was measured at 540 nm with a microplate reader (Synergy H1, BioTek, USA). Reducing sugar concentrations were calculated based on a calibration curve prepared with 0.2 – 2 g/L glucose.

### Backlit imaging of mycelium materials

Dried materials were placed on a drawing pad with white light (6500K; Crafts&Co, Netherlands) within a commercial lightbox (60×60cm; Puluz, China) for photography and photographed from above to capture backlit pictures (**Supplementary Figure S1**). Exposure was normalised within each image using a non-transmitting black reference as the minimum intensity value and the directly illuminated background as the maximum intensity value. Images were processed with a custom macro for ImageJ (version 1.54p; Schneider et al., 2012), with a mask enabling segmentation of the pellicle from the background. Transmittance was defined operationally as the natural logarithm of the mean grey value of the segmented pellicle.

### Scanning electron microscopy

Glycerol-treated mycelium pellicles were prepared for scanning electron microscopy (SEM) by cutting thin transversal slices using a scalpel to visualise the mycelial pellicle. Samples were mounted on aluminium SEM stubs using conductive carbon tape and gold-coated to a thickness of 3 nm using a Cressington 208HRD high resolution sputter Coater (Cressington Scientific Instruments, Watford,UK). SEM imaging was performed under vacuum using a JEOL JSM-6610LV SEM microscope equipped with an Oxford X-Max detector and Aztec software (Oxford Instruments, Germany). Images were acquired using secondary electron detection at an accelerating voltage of 5.0 kV, a probe current of 35.00 nA and a working distance ranging from 10 to 12 mm at magnifications ranging from 100- to 1000-fold.

### Tensile testing

Specimens were prepared from glycerol-treated, air-dried mycelium materials with a laser cutter (Glowforge Plus, Glowforge, WA, USA) at speed setting of 250 and power 60%–65% to match a dog-bone shape of 25 mm length, a gripping-end width of 10 mm, and a central gauge section 5 mm wide and 5 mm long (**Supplementary Figure S2**). Each biological replicate yielded between 5 and 9 specimens.

Tensile testing was performed on a Dynamic Mechanical Analyser (DMA Q800, TA Instruments, DE, USA). Tests were performed at room temperature. Width and thickness were measured using a calliper and thickness gauge, respectively. The test consisted of a constant force ramp of 3 N min^−1^, with a 0.01 N preload force and a maximum force of 18 N. Young’s modulus was calculated as the slope of the stress-strain curve in the linear elastic region between 1 and 10% strain. The highest stress value was considered the ultimate tensile strength (UTS).

### Statistical analysis and data visualisation

Data analysis and visualisation was performed in RStudio version 2025.09.2+418 using R version R 4.5.1 GUI 1.82 with the packages dplyr and ggplot2.

Reducing-sugar calibration curves were fitted by linear regression (absorbance against glucose concentration; 0.4–2.0 g L⁻¹; R² = 0.999). Concentrations below 0.8 g L⁻¹ were considered below the detection limit. Linear regression through the origin was used for the conversion of fresh to dry biomass (lyophilised against fresh biomass, n = 22). Biomass accumulation was modelled as a function of culture volume and incubation time using a second-order response surface model that included linear, quadratic and interaction terms (**Equation 1**). The predicted maximum was located by evaluating the fitted model over a grid of time and volume. Residual reducing-sugar concentration was modelled as a first-order linear function of incubation time and culture volume (**Equation 2**). For every volume, only time points up to and including the first measurement below 1 g L⁻¹ were included, thereby excluding points where reducing sugars were already depleted. As the interaction term did not improve the fit, only main effects were considered. Breaking force was regressed on specimen thickness for each bioprocess. Regression fits were evaluated using R² and overall model significance.

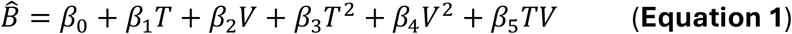

with *B̂* biomass accumulation, *T* incubation time and *V* culture volume.

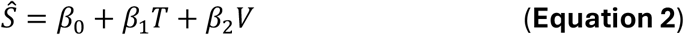

with *Ŝ* residual reducing-sugar concentration, *T* incubation time and *V* culture volume.

Descriptive statistics, including means, medians, and standard deviations, were calculated for each sample group. For non-normally distributed metrics, average values are reported as the median instead of the mean. Associations between continuous variables (transmittance and tensile strength, each with fresh biomass) were assessed with Pearson’s correlation coefficient. The relationship between breaking force and specimen thickness was assessed through linear regression and analysis of covariance (ANCOVA). Principal component analysis (PCA) was performed on centred and scaled metabolic, physical and mechanical variables, which were also used for PERMANOVA.

Statistical differences among groups were analysed through one-way or two-way ANOVAs when parametric assumptions were confirmed, followed by pairwise Tukey’s HSD post-hoc tests when significant differences were detected. If no transformation of the response variable resulted in satisfactory parametric assumptions, Kruskal-Wallis tests were used instead. When significant, pairwise Wilcoxon post-hoc tests with Benjamini–Hochberg correction were used. For multivariate comparisons, PERMANOVA with Holm correction applied to pairwise tests was used. Groups were considered statistically different when p < 0.05. Significance labels were assigned using compact letter displays or asterisks: ‘*’ for p < 0.05, ‘**’ for p < 0.01, and ‘***’ for p < 0.001.

## Results

### Liquid-state surface fermentation produces cohesive mycelium mats

In a first experiment, we assessed whether LSSF with *T. versicolor* could result in the formation of a cohesive material (**Figure 1**). The standard LSSF workflow consisted of a two-step inoculum preparation strategy (**Figure 1A**). A static liquid pre-culture of *T. versicolor* was blended, regenerated under shaken conditions and blended again to improve biomass dispersion. The resulting inoculum was added to fresh medium at a 1:5 inoculum-to-medium ratio (18 mL inoculum added to 90 mL medium) and incubated statically in the dark in round microboxes.

**Figure 1.**
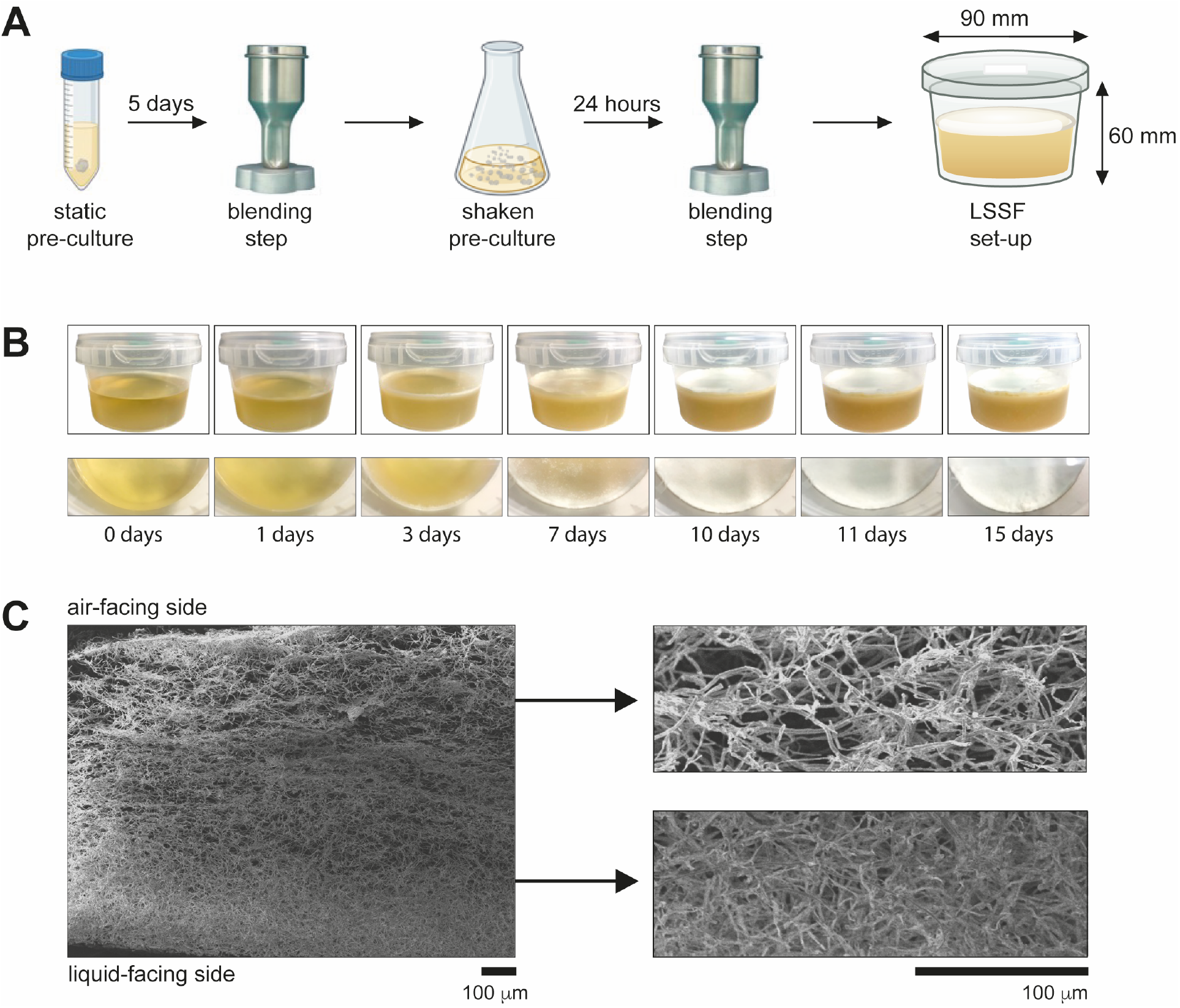
Fungal cultivation under liquid-state surface fermentation (LSSF). (**A**) Inoculum preparation: a static liquid pre-culture is blended, regenerated under shaken conditions, and blended again before inoculation of the production culture. (**B**) Progression of fungal growth under LSSF from day 0 to day 15; oblique (high-angle) view of the culture (top row) and top view of the developing pellicle (bottom row). (**C**) Scanning electron micrographs of a transverse section of a dried pellicle formed under LSSF (left), closeup views showing the air-facing and liquid-facing areas five times magnified (right). Scale bars: 100 µm (overview and insets). This are representative images for a larger set of SEM images.

Mycelium mat formation proceeded through distinct visually observable phases (**Figure 1B**). At the start of LSSF incubation, the inoculum was visible as translucent biomass dispersed throughout the liquid medium. During the first days of the static cultivation, turbidity of the liquid phase increased, pointing to fungal growth. By day 4, mycelial growth was observed at the air-liquid interface under the form of individual colonies. These developed into visible surface growth zones, which expanded radially over time. Growth zones gradually merged, and by day 6 the liquid surface was entirely covered by a continuous mycelial pellicle. Continued incubation until day 14 resulted in a thicker and more cohesive mat (**Figure 1B**). At that point, mycelium pellicles with a 90-mm diameter surface were harvested.

SEM imaging was used to examine the hyphal architecture of the harvested pellicles (**Figure 1C**). A side cross-section of a representative pellicle showed a clear vertical gradient in hyphal organisation, from the liquid-facing side to the air-facing side. The liquid-facing region showed a more compact hyphal network, whereas the air-facing region displayed a more open and porous structure, with a looser hyphal arrangement. The continuous interwoven hyphal network observed throughout the cross-section demonstrates a cohesive, stratified internal architecture of the LSSF-derived mycelium pellicle.

### Backlit imaging provides a readout of pellicle biomass and heterogeneity

To monitor mycelial development, mats were harvested at different incubation times and analysed by backlit imaging as a spatial readout of accumulated biomass (**Figure 2**). Dried mycelium pellicles were imaged on a light screen under standardised backlit conditions (**Supplementary Figure S1**); in this configuration light passes through the dried pellicle, so denser and thicker regions transmit less light. Backlit images (**Figure 2A**) were processed to segment the pellicle area, extract normalised greyscale values and express their mean as transmittance, defined here as its natural logarithm. Because light transmittance reflects pellicle thickness and hyphal density, it was used as an indirect proxy for biomass accumulation. Accordingly, transmittance decreased as biomass accumulated, tracking gravimetric biomass across incubation times (**Figure 2B**). Beyond the bulk estimate, backlit imaging also resolved the spatial distributed of biomass within the pellicle, distinguish regions of differing hyphal density and thickness (**Figure 2A**). Together, this shows that a single backlit image can both approximate bulk biomass and map its local variation across the mat.

**Figure 2.**
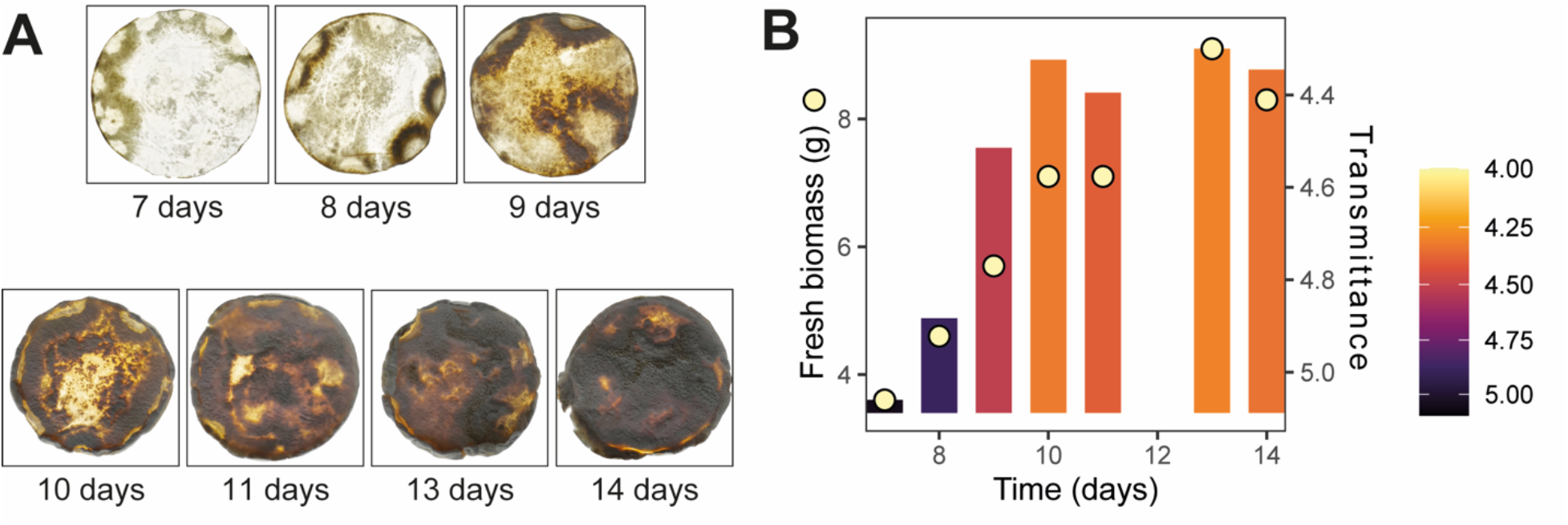
Backlit imaging of mycelium materials. (**A**) Backlit photographs of the dried biomass showing the increasing opacity of the material over incubation time, from day 7 to day 14. (**B**) Fresh biomass (points, left axis) and transmittance (bars, right axis; also encoded by bar colour) as a function of incubation time.

### Culture volume and incubation time determine the trajectory of biomass accumulation

To assess how culture volume and incubation time affect LSSF-mediated biomass accumulation, cultures were prepared at different liquid volumes and harvested after different incubation periods (**Supplementary Figure S3**), followed by biomass quantification using weighing and backlit imaging (**Figure 3**; **Supplementary Dataset S1**; **Supplementary Figure S4**). Culture volumes between 18 mL and 126 mL were incubated for 7, 8, 9, 10, 11, 13, and 14 days with the inoculum ratio being kept constant across all volumes at 1:5 (inoculum:medium, v/v).

**Figure 3.**
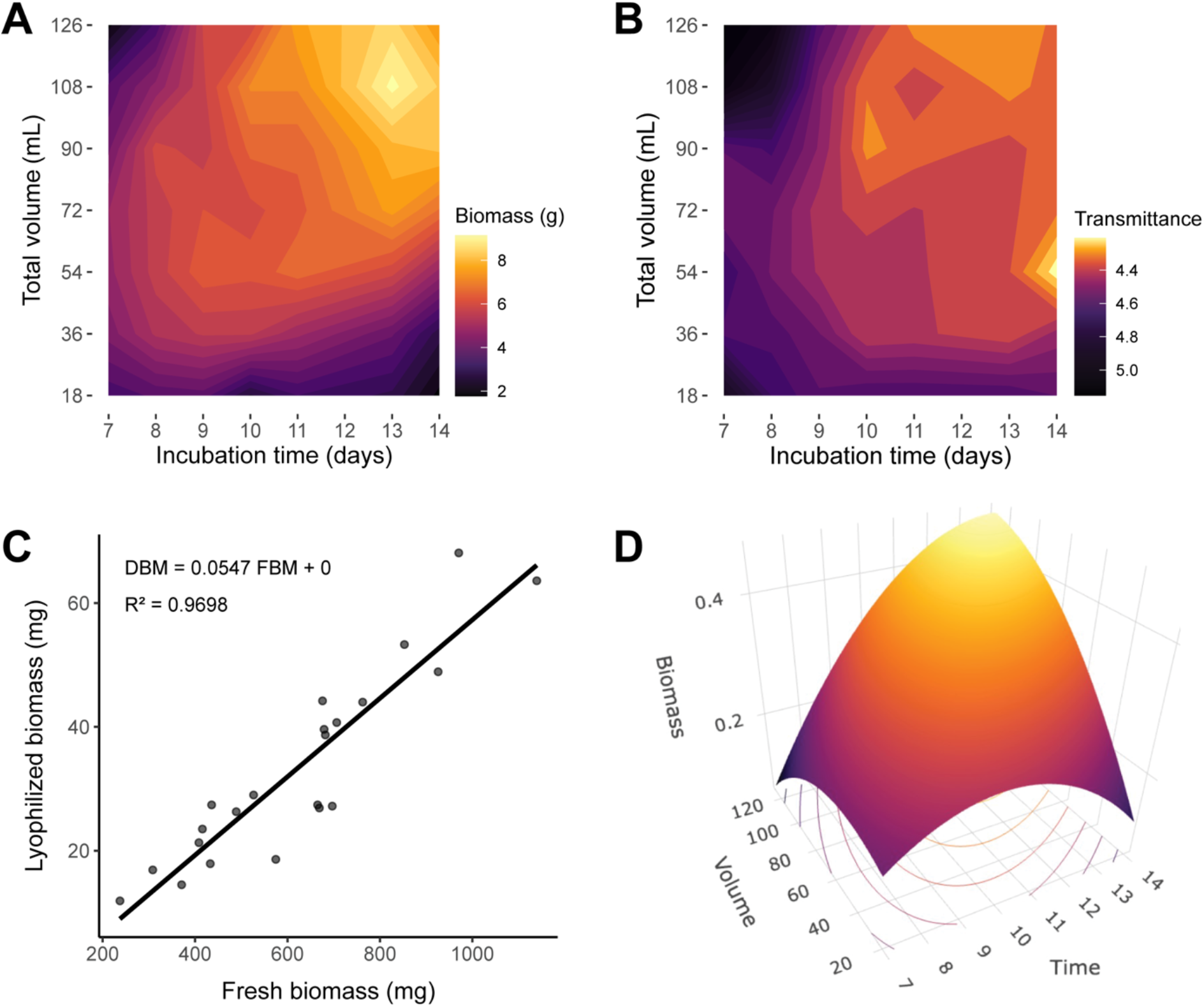
Impact of culture volume and incubation time on biomass accumulation under LSSF conditions. (**A**) Fresh biomass across different culture volumes (18 – 126 mL) and incubation times (7-14 days). (**B**) Light transmittance of biomass calculated as the natural logarithm of averaged grey values measured from backlit pictures of dried materials. (**C**) Linear regression of biomass weighed at harvest (fresh) and after vacuum-freeze drying (lyophilised; n = 22). (**D**) Second-order response surface model of biomass accumulation based on time, volume, their quadratic terms and interaction.

Culture volume influenced the timing and extent of surface colonisation and biomass accumulation (**Figure 3A**; **Supplementary Dataset S1**). Three volume-dependent phases could be distinguished: an initial biomass accumulation phase, a volume-specific phase in which the maximal biomass amount was obtained and, in some conditions, a subsequent biomass decline. Except for the 126-mL culture harvested on day 7, all tested conditions formed a continuous mycelial mat (**Supplementary Figure S3** and **S4**). Low-volume cultures (18 and 36 mL) formed continuous mats early, reaching a maximal biomass accumulation between 3.7 and 5.5 g around day 9, followed by a decline toward day 14 (**Figure 3A**). Intermediate volumes (54 and 72 mL) reached higher biomass values, with maxima between 6.6 and 7.5 g around days 11 and 13. In contrast, in high-volume cultures (90, 108 and 126 mL), biomass accumulation was delayed at early time points but continued until the end of the incubation period, reaching the highest values between 8.1 and 9.1 g (**Figure 3A**). Thus, lower culture volumes displayed an accelerated biomass accumulation but declined earlier, whereas larger volumes sustained biomass accumulation at later time points. The highest measured biomass was obtained after 13 days in a 108-mL culture.

Transmittance decreased as biomass weight measurements increased (**Figure 3B**; **Supplementary Dataset S1**). The two were strongly correlated over the full incubation (r = −0.7, p < 0.001, n = 49), strengthening to r = −0.87 (p < 0.001, n = 36) when restricted to the accumulation phase (time points up to peak biomass in each volume). This reflects a decoupling, where transmittance remained relatively stable until the end of incubation even where biomass declined (**Figure 3B**).

Because fresh biomass can vary with water content, biomass yields are conventionally reported in a dry-weight basis as a standard in biotechnology. To this end, dry weight was estimated from fresh weight using a fixed conversion established by a targeted validation experiment, in which fresh and lyophilised biomass were strongly correlated (R² = 0.97, p < 0.001, n = 22; **Figure 3**; **Supplementary Dataset S1**). The estimated dry weight data were then used to model biomass accumulation using a second-order response surface model as a function of incubation time, culture volume, their quadratic terms, and their interaction (**Figure 3D**):

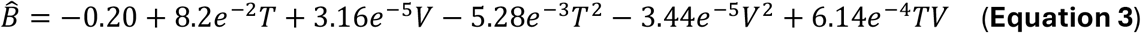

with *B̂* biomass accumulation, *T* incubation time and *V* culture volume.

The model provided a good fit to the experimental data, confirming that biomass accumulation was jointly shaped by incubation time and culture volume (**Figure 3D**; adjusted R² = 0.92, p < 0.001). Incubation time showed a significant positive linear term (*T*: β_1_ = 8.2 × 10^−2^, p < 0.001), while both quadratic terms were significant and negative (*T*²: β_3_ = −5.28 × 10^−3^, p < 0.001; *V*²: β_4_ = −3.44 × 10^−5^, p < 0.001), indicating a curved response surface.

In contrast, the linear volume term was not statistically significant (V: β_2_ = 3.16 × 10^−5^, p > 0.9). However, because the model includes quadratic and interaction terms, the non-significance of the linear volume term alone does not indicate that culture volume was unrelated to biomass accumulation. Indeed, a significant positive interaction between time and volume was detected (*TV*: β_5_ = 6.14 × 10^−4^, p < 0.001), consistent with the observed shift in biomass accumulation toward later incubation times at higher culture volumes. The fitted model predicted a theoretical maximum at an incubation time of 16 days and a culture volume of 146 mL, corresponding to approximately 0.48 g dry biomass (**Figure 3D**). This optimum lies just beyond the tested range (≤ 14 days, ≤ 126 mL) and should therefore be regarded as an extrapolation.

### Residual sugar concentration and pH reveal volume-dependent metabolic changes

To further assess the effects of culture volume and incubation time on LSSF performance, residual reducing-sugar concentrations were measured in spent media across the incubation period from 7 to 14 days (**Figure 4A**; **Supplementary Dataset S1**). Starting from an initial glucose concentration of 18.2 g L^−1^ (20 g L^−1^ of glucose monohydrate), residual reducing-sugar concentrations declined progressively for all culture volumes. This decline occurred earliest in low-volume cultures (18 and 36 mL), in which concentrations below 1 g L⁻¹ were already observed by day 8–9. In intermediate-volume cultures (54 and 72 mL), similarly low concentrations were reached later, between days 11 and 13. In the largest volumes (90–126 mL), residual reducing sugars remained detectable until the end of the incubation period (**Figure 4A**). The volume-dependent pattern of substrate depletion paralleled biomass accumulation trajectories (**Figure 3**): lower-volume cultures reached peak biomass earlier and subsequently declined in biomass, whereas higher-volume cultures still contained remaining reducing sugars in the spent medium and continued to accumulate biomass until the final sampling point (day 14).

**Figure 4.**
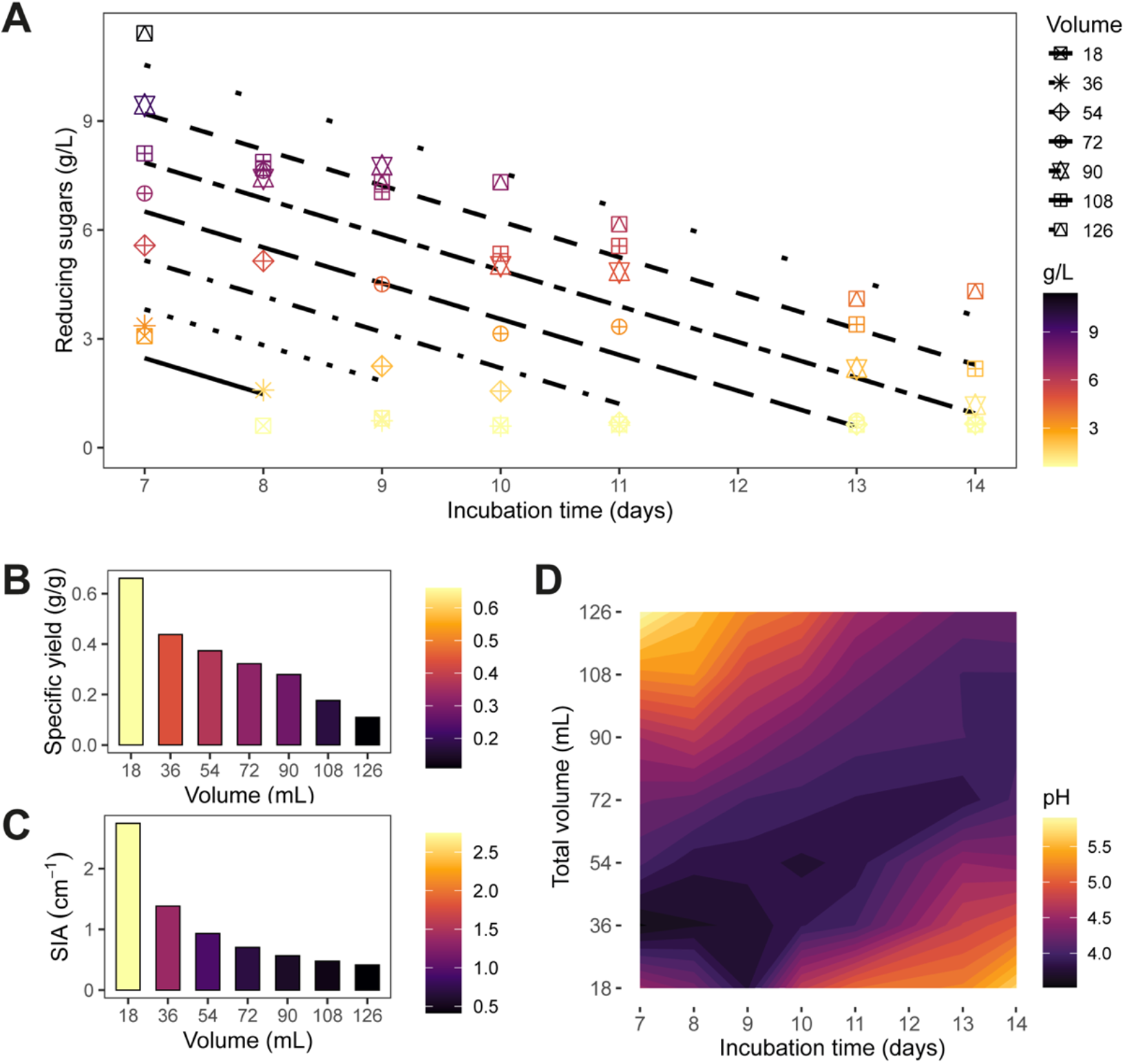
Effects of culture volume, incubation time and specific interfacial area (SIA) on substrate utilisation and medium pH during LSSF. **(A)** Residual reducing-sugar concentration in spent medium as a function of incubation time and culture volume. Points represent the average of three technical replicates, coloured according to reducing-sugar concentration in spent medium as a function of incubation time. Lines represent predictions from the linear model including incubation time and culture volume as main effects, without an interaction term. Different line types correspond to different culture volumes**. (B)** Maximum specific biomass yield obtained for each culture volume after 7 days incubation (Supplementary Dataset S1). **(C)** Specific interfacial area (SIA; surface-to-volume ratio; Supplementary Dataset S2). **(D)** Contour plot of spent-medium pH at the harvesting moment as a function of incubation time and culture volume.

Substrate consumption was modelled over the active-consumption phase. For each culture volume, this comprised all time points up to and including the first measurement below 1 g L⁻¹, thus excluding points where reducing sugars were already depleted. Later samples represented the near-depletion phase, which coincided with the onset of biomass decline in low-volume cultures (**Figure 3A**). A first-order linear model of residual reducing-sugar concentrations was developed as a function of incubation time and culture volume (**Figure 4A**):

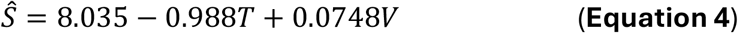

with *Ŝ* the remaining glucose in the spent media, *T* the incubation time and *V* the culture volume.

The model provided a good fit to the data (adjusted R² = 0.89, p < 0.001). In contrast to the model for biomass accumulation, the interaction term between time and volume did not improve the model and was therefore not retained. Incubation time was negatively associated with residual reducing-sugar concentration (*T*: β_1_ = −0.99, p < 0.001). Sugar concentration decreased linearly at a constant rate of approximately 1.0 g L⁻¹ d⁻¹. In contrast, culture volume was positively associated with residual reducing-sugar concentration (V: β_2_ = 0.075, p < 0.001). These volume-dependent dynamics paralleled that of biomass accumulation. Low-volume cultures depleted reducing sugars earlier, reached a maximal biomass peak sooner, after which biomass declined. In contrast, availability of reducing sugars was retained for a longer time in higher-volume cultures, enabling biomass accumulation until the final sampling point. Finally, because the interaction term was not retained, the substrate depletion model predicted parallel linear declines in residual reducing-sugar concentration over time across culture volumes (**Figure 4A**).

Biomass accumulation and reducing-sugar depletion data were used to calculate fermentation performance parameters across culture volumes (**Supplementary Dataset S1**; **Supplementary Text S1**). This revealed a trade-off between absolute biomass and substrate-use efficiency. Although larger culture volumes produced more biomass overall, substrate was converted to biomass more efficiently at lower volumes, particularly at early incubation times (**Supplementary Dataset S1** and **S2**; **Figure 4B** and **4C**). The highest biomass yield on substrate (Y_x/s_) was observed after 7 days in the 18-mL culture (0.66 g g^−1^) and decreased progressively with increasing culture volume, reaching 0.11 g g^−1^ at 126 mL (**Figure 4B**). This could be explained by a higher specific interfacial area (SIA; interfacial surface area per culture volume) at lower culture volumes (**Figure 4C**).

LSSF incubation was also associated with volume- and time-dependent pH decreases (**Figure 4D**; **Supplementary Dataset S1**). Low-volume cultures displayed the strongest and earliest acidification, reaching pH values below 4 by day 9, followed by an increase at later time points. In contrast, pH declined more gradually in larger-volume cultures and approached approximately pH 4 later in incubation. The stronger pH decrease in low-volume cultures coincided with their higher specific biomass yields (**Supplementary Dataset S1**), consistent with a higher metabolic activity and fermentation performance. The subsequent pH increase in low-volume cultures occurred when residual reducing-sugar concentrations were low and biomass declined, suggesting reduced metabolic activity during the final incubation phase.

### Inoculum source and processing determine biomass yield

To assess the effect of inoculum preparation on LSSF-derived mycelial pellicles, different approaches were compared (**Figure 5**). Inocula from agar plates (A) or static liquid cultures (L) were subjected to different processing steps: one blending step (B), blending followed by shaken incubation (BS) or blending-shaking-blending (BSB) (**Figure 5A**). Each inoculum was added to 90 mL of fresh medium and incubated statically for 14 days. Harvested pellicles were evaluated for biomass amount (**Figure 5B**) and heterogeneity by backlit imaging (**Supplementary Figure S5**).

**Figure 5.**
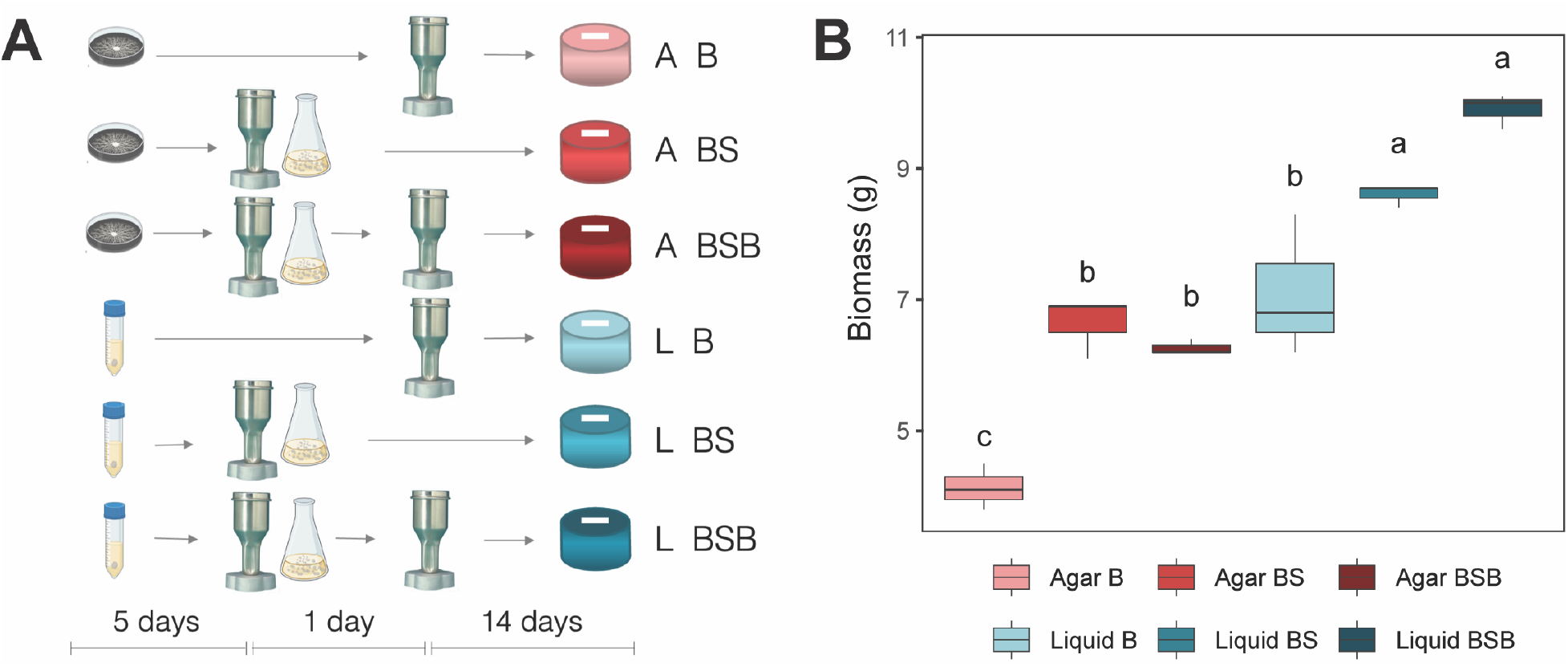
Inoculum source and processing influence LSSF biomass yield. (**A**) Experimental design comparing two biomass sources, agar plate (A) and liquid static culture (L), and three inoculum-preparation protocols: one blending step (B), blending followed by shaken incubation (BS), and blending-shaking-blending (BSB). (**B**) Fresh biomass produced after 14 days of LSSF using the six inoculum types. Different letters indicate statistically significant differences between groups, whereas groups sharing a letter are not significantly different (α = 0.05; n = 4).

Cultures inoculated with biomass derived from static liquid culture produced significantly higher biomass than those initiated from agar plates (**Figure 5B**; mean values of 8.5 g and 5.7 g, respectively; p < 0.001). Inoculum-processing steps also affected biomass yield across both biomass sources. The shaken-incubation protocols (BS and BSB) resulted in higher biomass accumulation (7.6 g and 8.1 g, respectively), whereas the single-blending process (B) produced the lowest yield (5.6 g; p < 0.001). Thus, static liquid-culture-derived inocula and the inclusion of a shaken incubation step increased biomass yield during LSSF. The second blending step did not significantly further alter biomass yield relative to BS (**Figure 5B**).

### Inoculum processing affects mechanical properties of pellicles

Because LSSF directly produces coherent mycelium pellicles, their mechanical properties were assessed by tensile testing (**Figure 6; Supplementary Table S1**). After glycerol treatment and air-drying, dog-bone-shaped specimens were laser-cut from the pellicles for tensile testing (**Figure 6A**; **Supplementary Figure S2**). The tensile behaviour of the specimens was captured in stress-strain curves (**Figure 6B**) with the ultimate tensile strength representing the maximum stress reached during each test (**Figure 6C**).

**Figure 6.**
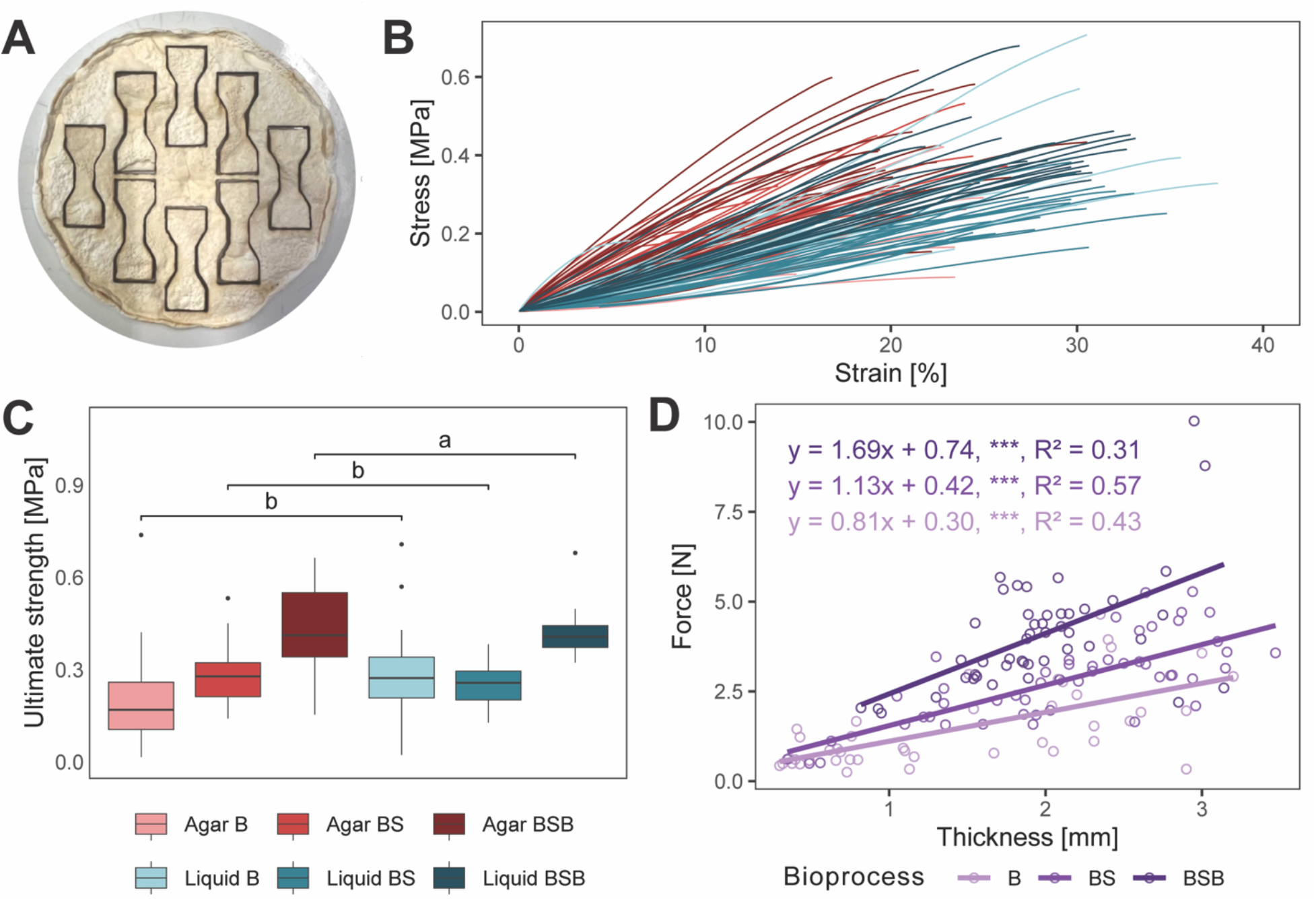
Mechanical characterisation of pellicles produced with different inoculum preparation strategies. (**A**) Layout of dog-bone specimens laser-cut from a pellicle. (**B**) Stress-strain curves of individual specimens for the six inoculum types, which combine two biomass sources with three bioprocesses (n ≥ 18 specimens per inoculum type). (**C**) Ultimate tensile strength. Boxes show the median and interquartile range (IQR); whiskers extend to 1.5 × IQR. Brackets group inoculum types sharing the same bioprocess. Different letters indicate significant differences between bioprocesses (Kruskal–Wallis test followed by pairwise Wilcoxon rank-sum tests with Benjamini–Hochberg correction, p < 0.05). (**D**) Relationship between maximum force and pellicle thickness per bioprocess. Lines show linear regression fits, with the equation, encoded p-value and R^2^. Bioprocess abbreviations as in Figure 5.

Pellicles produced using static liquid culture-derived and agar-derived inocula had similar median ultimate tensile strengths (0.33 MPa and 0.29 MPa, respectively) (**Figure 6C**). However, tensile strength varied with the inoculum-processing protocols. A single blending step yielded the lowest median strength (B = 0.24 MPa). When a subsequent shaken incubation step was included, the median strength increased (BS = 0.28 MPa; p < 0.05). The strongest materials resulted from a second blending after the shaken incubation (BSB = 0.41 MPa; p < 0.001).

Breaking force scales with the specimen cross-section, so raw force confounds strength with thickness. Plotting force against thickness separates them (**Figure 6D**). At matched thickness, the more elaborate processes sustained a higher force (p < 0.001), and the force gained per unit thickness also increased with processing (p < 0.05). Across all specimens, ultimate tensile strength was only weakly correlated with fresh biomass (r = 0.33, p < 0.001).

Despite similar median tensile strengths, inoculum source affected pellicle geometry and deformation (**Supplementary Table S1**). Compared with liquid-inoculated pellicles, agar-inoculated ones were thicker (2.09 vs. 1.77 mm; p < 0.01) and had a larger surface area after drying (45.8 cm² vs. 38.7 cm²; p < 0.001), a lower strain at break (19.3% vs. 27.5%; p < 0.001) and a higher Young’s modulus (1.68 MPa vs. 1.18 MPa; p < 0.001). Including a shaken incubation step (BS) yielded more biomass and thicker, slightly stronger and more stretchable pellicles than a single blending step (B) (Supplementary Table S1). Because tensile strength and strain at break increased together, Young’s modulus remained unchanged. A second blending step (BSB) further increased thickness and considerably increased tensile strength without changing strain at break or biomass, resulting in a higher Young’s modulus (**Supplementary Table S1**).

Integrating all measured traits (metabolic, physical and mechanical), principal component analysis separated the six inoculation strategies, the first two components explaining 61.8 % of the variance (**Supplementary Figure S6**). PERMANOVA attributed 64.2 % of the multivariate variance to treatment, and all groups differed significantly (p < 0.05). Each inoculation strategy thus produced a distinct multivariate signature. Biomass source separated samples mainly along PC1, whereas inoculum processing separated them along PC2, with B, BS and BSB ordered by increasing processing.

## Discussion

Pellicle formation within LSSF proceeds through a sequence of physical and physiological stages that can be mapped onto the traditional microbial growth phases. The fungus first grows submerged in the static liquid medium, where oxygen diffusion limits growth and biomass accumulation as well as substrate consumption are slow. Analogous to the microbial lag phase, this stage represents the delay before surface colonisation. Escape to the air-liquid interface is facilitated by hydrophobins, which lower water surface tension (Wösten and Willey, 2000), though the process remains mechanistically unclear. Once at the air-liquid interface, where oxygen is no longer limiting, biomass accumulation and nutrient uptake occur rapidly, resembling the exponential growth phase. Colonisation of the interface begins at multiple points and times across the surface, each acting as a nucleation site for radial expansion. Neighbouring colonies coalesce until the interface is fully covered, constrained by vessel geometry. The timing, density and distribution of these nucleation events shape the homogeneity and properties of the pellicle. Once coverage is complete, substrate consumption becomes steady and biomass accumulates slowly as the pellicle thickens and densifies, consistent with the stationary phase. Throughout, metabolic activity acidifies the medium as low as pH = 3.7 via accumulation of organic acids and dissolved CO_2_ (Liaud *et al*., 2014). Biomass decline is better understood as a consequence of overgrowth than as a growth phase: once the pellicle has outgrown its carbon supply, glucose depletion leaves it unable to sustain itself, the acidification reverses, and biomass declines.

In static liquid cultures, oxygen is supplied by diffusion from the surface, so the specific interfacial area (SIA; aerial surface area per culture volume) sets mass-transfer conditions. In static kombucha fermentation, equal SIA gives comparable fermentation efficiency regardless of absolute volume (Malbaša *et al*., 2006; Cvetković *et al*., 2008). Similarly, in liquid static cultures of Ganoderma lucidum, ganoderic acid production decreased as culture volume increased, which was attributed to reduced oxygen supply to the surface mat (Hu *et al*., 2018). By analogy, a higher SIA in this system means a shallower, better-oxygenated column and a shorter lag (2.75 to 0.41 cm⁻¹, lowest to largest volume; **Figure 4C**). Consistent with this, mean glucose uptake per litre over the first 7 days intensified in lower volumes (from 2.2 g L⁻¹ d⁻¹ at 18 mL to 1.0 g L⁻¹ d⁻¹ at 126 mL), so that the smallest volumes had consumed over 80 % of their glucose by day 7, compared with 37 % at 126 mL (**Supplementary Dataset S2**). Absolute uptake, however, increased with culture volume (0.27 to 0.85 g per vessel) even though no continuous mat had yet formed at 126 mL (**Supplementary Figure S4**). A large proportion of glucose was therefore consumed below the interface, likely by respiration and by submerged growth, which tracks with the inoculum added at a fixed volumetric ratio and competes for nutrients with pellicle formation.

Pellicle densification, in contrast, is limited by nutrient availability per unit interfacial area. After day 7, uptake converged to a constant volumetric rate across volumes (∼1.0 g L⁻¹ d⁻¹), so uptake per unit interface increased with volume (3.6 to 24 g m⁻² d⁻¹; **Supplementary Dataset S2**). At that point, the glucose remaining per unit interface differed 25-fold between volumes (11 to 277 g m⁻²) and tracked subsequent densification: pellicles with less than ∼60 g m⁻² remaining stopped growing or lost biomass between days 7 and 14, whereas larger reservoirs supported further gains of 21 to 59 g m⁻² dry biomass (Supplementary Dataset S2). The glucose load per unit area (66 to 442 g m⁻²) thus sets an upper limit to mat thickening.

Lag and densification thereby respond oppositely to culture volume. Reducing volume shortens lag and raises substrate-use efficiency but limits densification and absolute biomass. The highest yield (0.66 g g⁻¹ dry basis, 18 mL; **Figure 4**) lies at the upper end of biomass yields reported for filamentous fungi on glucose (0.25–0.67 g g⁻¹ in *Penicillium*; (Grijseels *et al*., 2017). Large volumes densify more but colonise later, and their efficiency is lower partly because glucose consumed by submerged biomass is not recovered in the pellicle. As SIA rises with 1/V, efficiency approaches a ceiling at high SIA, where the limit shifts from oxygen supply to the intrinsic metabolic efficiency of the fungus. Strategies that shorten lag while extending densification, such as a higher substrate concentration or a nutrient-rich agar layer at the vessel base, could combine early colonisation with sustained thickening.

Backlit imaging provided a non-destructive, image-based readout of pellicle biomass, conceptually analogous to optical-density measurements in suspension (Posch *et al*., 2012; Carrillo-Acuña *et al*., 2023). Quantitatively, the whole-pellicle mean transmittance estimated total biomass (**Figure 2**; **Supplementary Figure S4**); qualitatively, the image itself revealed how biomass was distributed, exposing heterogeneity and spatial patterns across the mat (Figure 2A). Because the method preserves the full two-dimensional transmittance map, these observations could be quantified further. For example, sub-region means could provide localised, non-destructive biomass estimates, transmittance variability (like variance or skewness) could measure heterogeneity, and analysis of adjacent regions could enable pattern or texture characterisation of a mat’s mesostructure.

Biomass yield depended on both inoculum source and processing. Liquid pre-cultures outperformed agar plates. Agar resists mechanical disruption, giving poorer dispersion, and the shift from solid to submerged growth requires physiological adaptation that delays growth. Accordingly, a shaken pre-culture step raised biomass (BS vs B), likely by easing this adaptation and shortening the lag. A second blending step (BSB vs BS) left biomass unchanged, suggesting the shaken-incubation pellets were already well dispersed and unaffected by further blending.

Alongside biomass, the shaken step (BS) raised tensile strength, consistent with earlier pellicle maturation and a longer densification period, and the second blend (BSB) raised it further (**Figure 6C**). Because BS and BSB reached similar biomass (**Figure 5B**), and breaking force increased at matched thickness (**Figure 6D**), this further gain is not explained by more biomass, but rather reflects an intrinsic material property. Consistent with this, strength was only weakly correlated with biomass across all samples (r = 0.33, p < 0.001). Additional fragmentation improved inoculum dispersion (Liu *et al*., 2016), reducing local heterogeneity in the BSB pellicles (consistent with the uniform appearance of their backlit images; **Supplementary Figure S5**) and thus the weak points from which fracture initiates and propagates.

The highest median tensile strength obtained here was 0.41 MPa (BSB), from coherent mats harvested directly from the culture, without filtration, pressing or other post-processing. Reported values for PMM range from 0.27 to 7.8 MPa across LSSF (0.27 to 1.5 MPa; César et al., 2021; Elsacker et al., 2023), SSSF (0.8 to 1.1 MPa; Haneef et al., 2017) and SmF, where biomass is typically blended, filtered and dried into sheets (3.7 to 7.8 MPa; (Appels *et al*., 2020; Wijayarathna *et al*., 2022; d’Errico *et al*., 2025; Jo *et al*., 2025; Van Den Brandhof *et al*., 2025). These benchmarks point to three levers for improving the mechanical performance of LSSF materials. First, harvest timing: the glucose consumption dynamics observed here show that the tested pellicles were approaching substrate depletion, and strength has been reported to decline with prolonged incubation (Elsacker *et al*., 2023). Harvesting closer to peak biomass should yield stronger materials. Second, cultivation conditions: in floating mats from *S. commune*, light and CO_2_ levels altered tensile strength up to 2.6-fold, reaching 40 MPa in a hydrophobin-deficient strain (Appels *et al*., 2018). In this study, inoculum preparation had a comparable effect: BSB raised strength 1.7-fold relative to B, consistent with a more homogeneous network rather than more biomass. Third, strain selection: mechanical performance can vary widely even within a single species under identical cultivation and processing (d’Errico *et al*., 2025). Between species, van den Brandhof et al., (2025) found no significant differences in PMM produced by SmF, possibly because blending and filtration during postprocessing mask native, species-specific network traits. Under LSSF, where mycelium pellicles remain intact, César et al., (2021) obtained values comparable to ours, and their data hint that species with ligneous (woody) fruiting bodies, such as *Ganoderma* spp., form stronger pellicles than species with coriaceous (leathery) ones. This would account for the higher strength reported by Elsacker et al., (2023) with a ligneous species.

Agar-derived pellicles were thicker and slightly larger than liquid-derived ones despite lower biomass, and thus less dense (**Supplementary Table S1**). Such differences presumably stemmed from agar fragments incorporated into the pellicle, which may also have limited shrinkage during drying. At comparable ultimate tensile strength, the thicker agar-derived pellicles required a higher force to break. Their lower strain at break is consistent with agar acting as a rigid filler that restricts network rearrangement under load (Fu *et al*., 2008). A shaken incubation step increased stretchability, plausibly by diluting the agar carried into the final culture and by supplying more malleable biomass than plate-derived inoculum, whereas the second blending step left it unchanged.

Considering all measured traits together (metabolic, physical and mechanical), each strategy produced its own distinct signature (**Supplementary Figure S6**). Biomass source and processing mapped onto separate principal components, indicating that they act on largely distinct sets of traits and could therefore be tuned separately. Processing behaved as a graded lever, with each additional step shifting the material further along the same component (**Supplementary Figure S6**).

Together, the two experiments define practical design rules for LSSF. Culture volume imposes a trade-off: shallow, high-SIA cultures convert substrate more efficiently, whereas deeper ones yield more biomass at lower efficiency. A more effective approach is to decouple the two, for instance by increasing substrate concentration in shallow cultures or partitioning larger volumes into shallow compartments. Harvest is best timed to peak biomass, before carbon depletion triggers decay. *In situ* monitoring of biomass accumulation through backlit imaging could provide a continuous readout to time harvest. More elaborate inoculum processing, such as shaking and blending, yields more homogeneous and stronger pellicles. Requiring neither agitation, aeration nor a solid substrate, and yielding coherent mats without post-processing, LSSF offers a promising low-energy route to PMM.

## Conclusions

Here, we propose liquid-state surface fermentation (LSSF) as a promising approach to the production of pure mycelium materials (PMM) with minimal energy input and without the labour-intensive practices typically involved in other fermentation approaches. We show the process of LSSF to produce PMM with *Trametes versicolor*, starting from inoculum preparation, analysing growth dynamics and glucose consumption until biomass harvest.

This study proposes a mechanism for interfacial pellicle formation divided into three different phases. The initial step is characterised by growth in suspension, limited by oxygen diffusion and therefore favoured by a higher specific interfacial area (SIA). After surface colonisation, rapid radial expansion takes place on the interfacial surface, constituting the second phase. Once the pellicle has reached full coverage, densification takes place in the third phase, restricted by glucose availability per unit interface. During this phase, the culture consumed glucose at a common volumetric rate (∼1.0 g L⁻¹ d⁻¹) across volumes, whereas much of the glucose consumed earlier by deeper cultures was not recovered in the pellicle, indicating competition from submerged growth. Smaller volumes thus raised the yield on glucose (up to 0.66 g g⁻¹) but limited densification and absolute biomass**, and biomass accumulation reversed once glucose was depleted.

We also demonstrated the relevance of specific choices in the process of inoculum preparation for material performance, including mechanical characterisation of the mycelium pellicle. Both the use of biomass derived from static liquid culture and the inclusion of a shaken liquid incubation step during inoculum preparation resulted in higher biomass. Furthermore, a second blending step further increased tensile strength without additional biomass, consistent with a more homogeneous pellicle. Additionally, agar-initiated cultures produced thicker, less dense pellicles that behave mechanically as particle-filled mycelial composites: stiffer and less stretchable yet maintaining comparable tensile strength. Backlit transmittance imaging provided a non-destructive estimate of pellicle biomass and its spatial distribution.

Altogether, these findings inform the production of mycelium sheets, including inoculum preparation and process conditions such as SIA and glucose load per unit interface. This work provides a basis for LSSF technologies and process-level design rules for PMM production.

## Authors contribution

**Jara Saluena Martin**: conceptualization, data curation, formal analysis, investigation, methodology, software, visualization, writing – original draft preparation, writing – review & editing. **Piet Dimitriadis**: investigation, software, writing – review & editing. **Soufian Attalibi**: data curation, investigation, visualization. **Annah-Ololade Sangosanya**: investigation, writing – review & editing. **Rafael Philippini**: investigation, methodology, supervision, writing – review & editing. **Razieh Shafiei-Alavijeh**: supervision, writing – review & editing. **Joost Brancart**: supervision, writing – review & editing. **Elise Elsacker**: conceptualization, supervision, writing – review & editing. **Eveline Peeters**: conceptualization, funding acquisition, supervision, writing – review & editing.

## Acknowledgments

The authors would like to thank Anouk Verstuyft for producing the 3D renders used in the illustrations of this study.

## Data Availability Statement

All data underlying this study are available in Supplementary materials and on Zenodo (10.5281/zenodo.23433712). All code scripts utilised during this work are available in Github (https://github.com/JaraTech/LSSF_data_analysis).

