## Supplementary Material for "Liquid-state surface fermentation as a platform for pure mycelium materials production"

Saluena Martin *et al.*

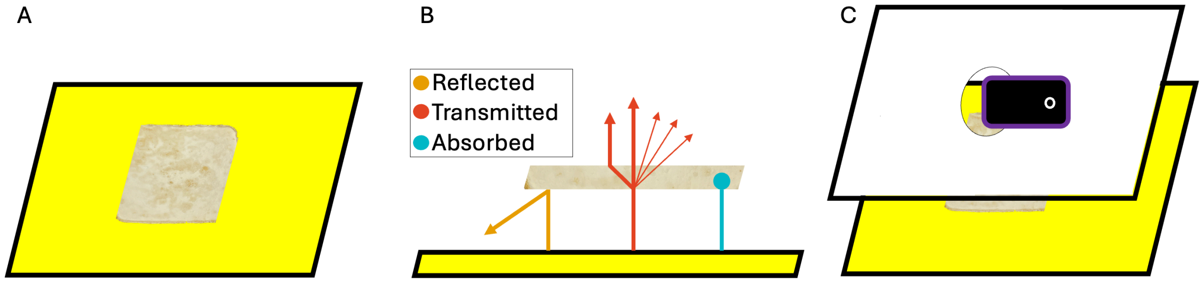


**Supplementary Figure S1.** Setup to acquire backlit images of mycelium pellicles. (**A**) mycelium pellicle is laid on a white light screen. (**B**) When encountering the mycelium pellicle, light might be reflected (bouncing off the pellicle) absorbed by the material or transmitted, comprising processes like scattering, refraction or diffraction. (**C**) the image was acquired at a constant distance from the object.

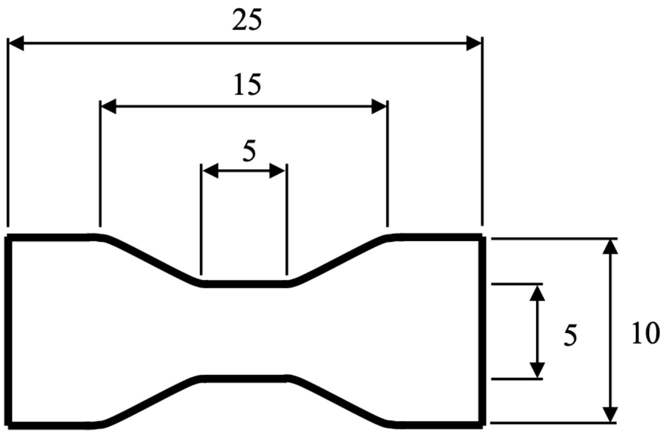


**Supplementary Figure S2.** Generation of specimens for a tensile testing. Dog-bone shape of 25 mm length, a gripping-end width of 10 mm, and a central gauge section 5 mm wide and 5 mm long.

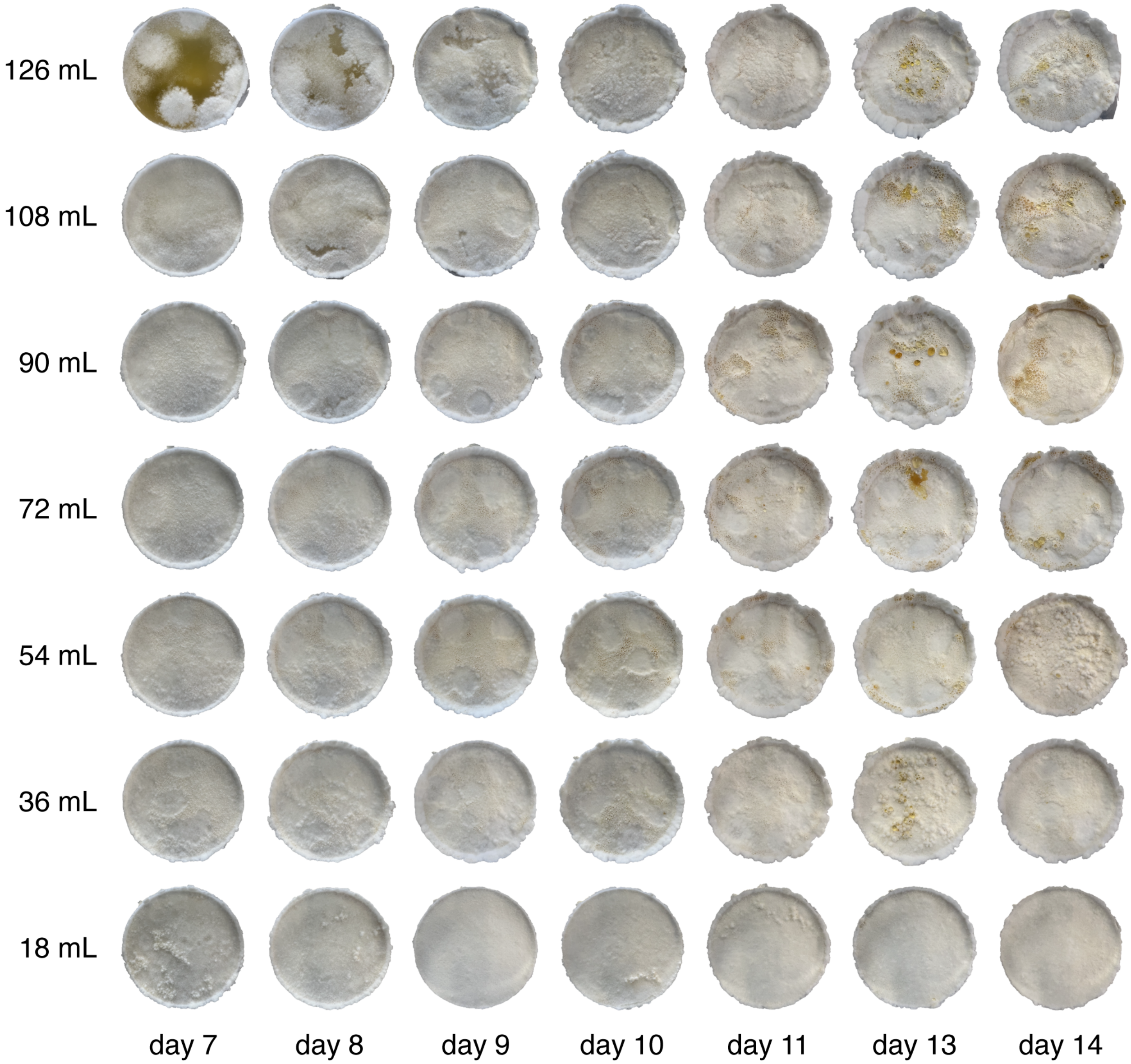


**Supplementary Figure S3**. Pictures of LSSF culture vessels across different culture volumes (18 – 126 mL) and incubation times (7-14 days).

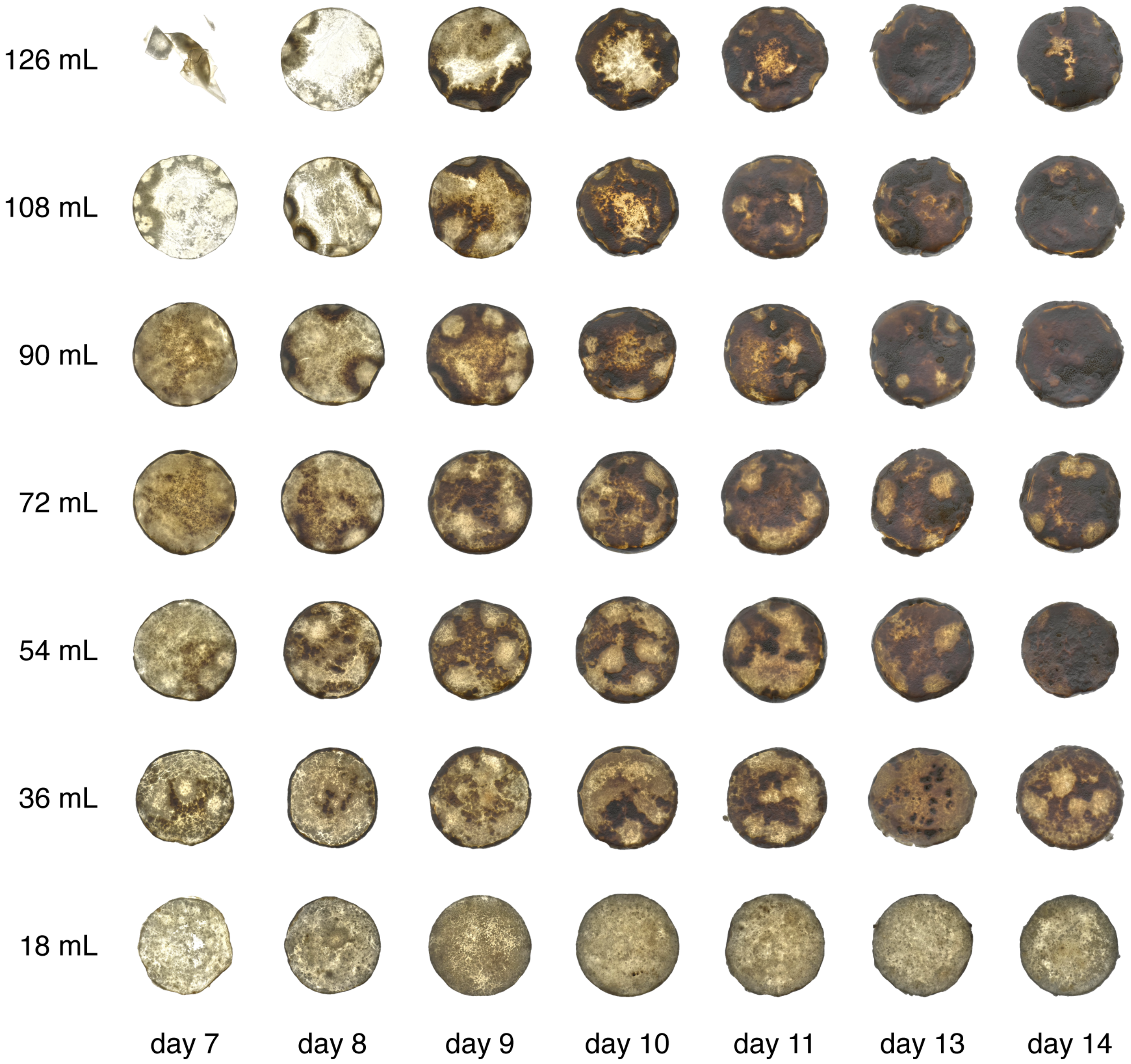


**Supplementary Figure S4.** Backlit pictures of air-dried mycelium pellicles grown under LSSF across different culture volumes (18 – 126 mL) and incubation times (7-14 days).

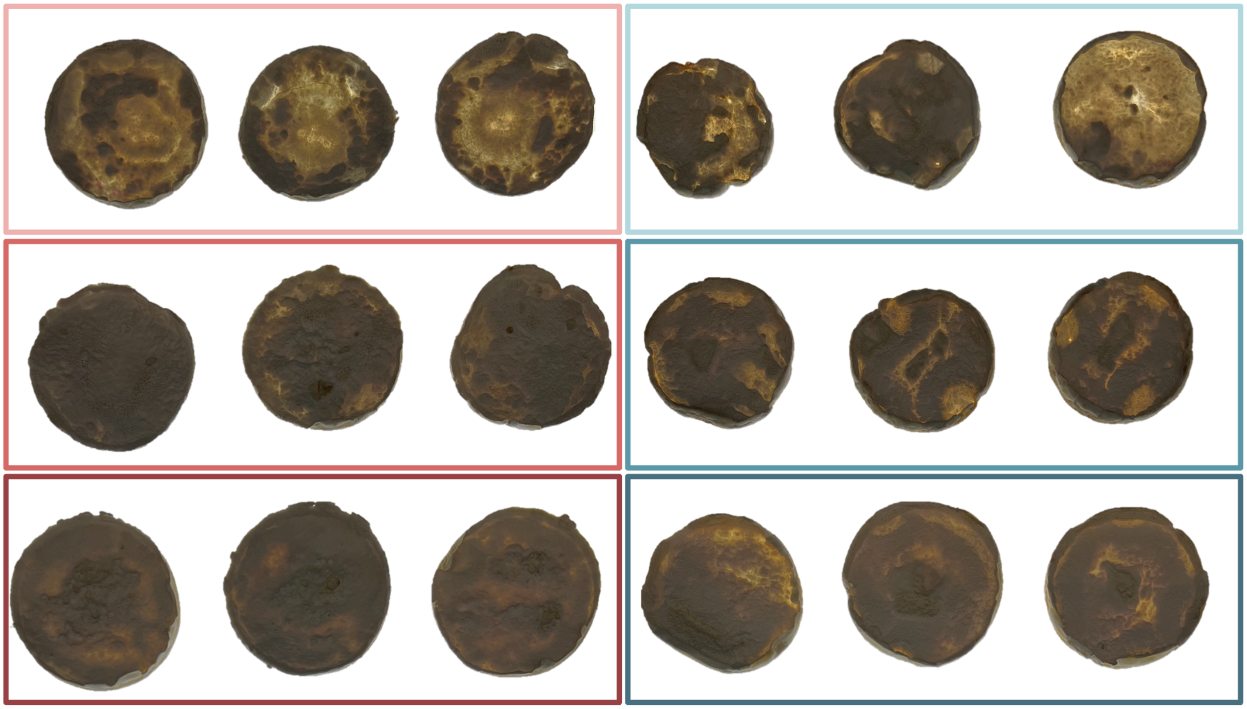

**Supplementary Figure S5.** Backlit pictures of biomass produced following different inoculum preparation bioprocesses. Two different biomass sources, agar plate (red hue) and liquid static culture (blue hue), and three different bioprocesses involving one (light shade) or two blending events with (dark shade) or without a shaken step in between (intermediate shade).

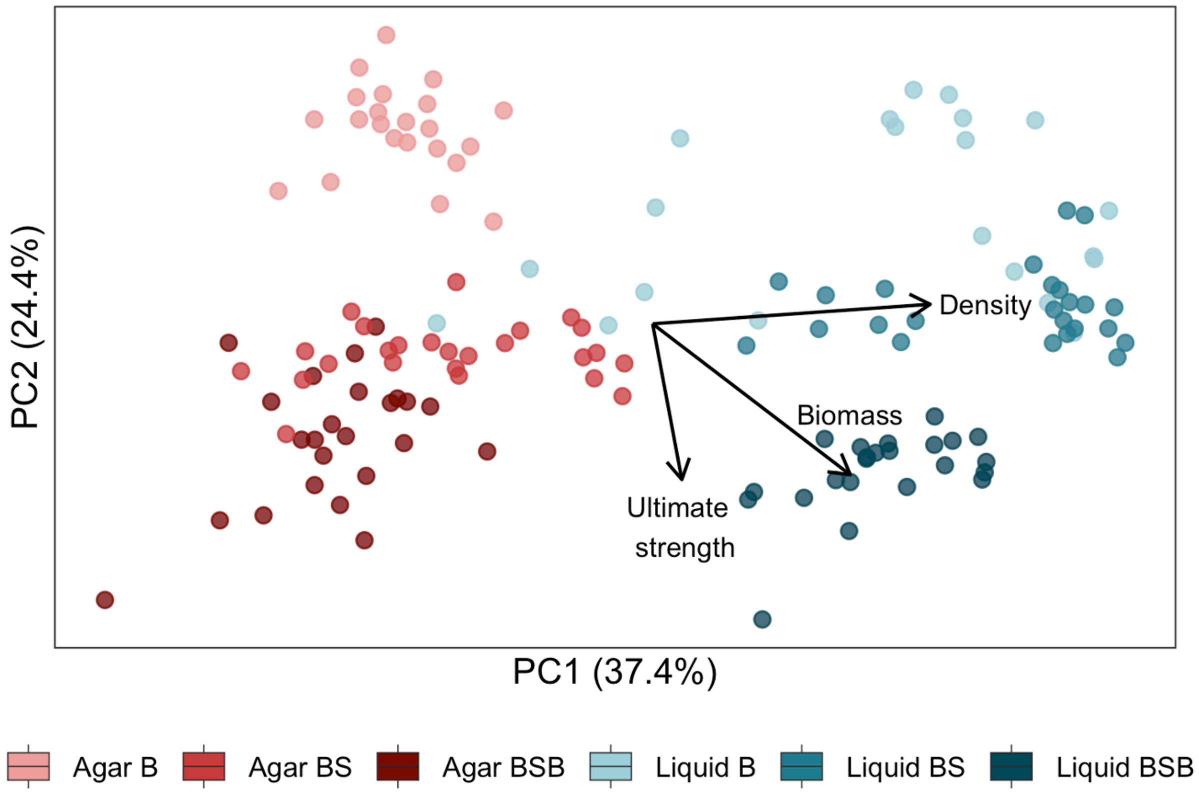


**Supplementary Figure S6**. Biplot of Principal Component Analysis (PCA) showing PC1 and PC2 of mycelium pellicles using different strategies for inoculum preparation: two different biomass sources (agar plate and liquid static culture), and three different bioprocesses involving one blending event (B) plus a shaken incubation step (BS) plus a second blending event (BSB). Parameters accounted for in the PCA are shown in Supplementary Table S3. Eigenvectors of pellicle density, treated dry biomass, and ultimate strength are represented with an arrow.

**Supplementary Table S1**. Data from pellicles produced using different strategies for inoculum preparation. Data grouped by biomass source (agar and liquid), bioprocess (B, BS and BSB) and by the combination of both (A-B, A-BS, A-BSB, L-B, L-BS, and L-BSB). pH indicates the pH of spent media at the time of pellicle harvest. Dry biomass shows the calculation of biomass based on fresh quantification at harvest and regression equation in **Supplementary Figure S3**. Glycerol weight represents the weight of glycerol treated and air-dried pellicles. Pellicle surface reflects the pixel area converted into mm$^2$ measured from backlit pictures of air-dried pellicles. The following column (N) indicates the number of replicates in each group. At least 5 specimens for further tests were laser cut from each biological replicate. Pellicle thickness was extracted from the thickness of each tensile specimen. Force at break displays the force required to break the material, irrespective to thickness. Ultimate strength shows the stress at specimen failure. Strain indicates the percentage of stretch when the specimen broke. Young's modulus was calculated from the tensile data between 1 – 10 % strain.

| | pH | Fresh biomass (g) | Glycerol biomass (g) | Pellicle surface (mm$^2$) | Apparent density (kg/m$^3$) | N | Pellicle thickness (mm) | Force at break (N) | Ultimate tensile strength (MPa) | Strain at break (%) | Young's modulus (MPa) | N |
|---|---|---|---|---|---|---|---|---|---|---|---|---|
| Agar B | 4.22±0.03$^{ab}$ | 4.13±0.35$^{c}$ | 0.76±0.03$^{c}$ | 44.1±2.93$^{a}$ | 128.7±15.33$^{a}$ | 3 | 1.1±0.73$^{c}$ | 0.86±0.63$^{c}$ | 0.17±0.10$^{c}$ | 15.2±5.50$^{c}$ | 0.87±0.58$^{de}$ | 21 |
| Agar BS | 4.09±0.04$^{cd}$ | 6.63±0.46$^{b}$ | 1.28±0.07$^{b}$ | 44.4±1.77$^{a}$ | 116.0±24.5$^{a}$ | 3 | 2.62±0.56$^{a}$ | 3.37±0.85$^{a}$ | 0.29±0.09$^{b}$ | 19.3±4.98$^{c}$ | 1.63±0.47$^{b}$ | 22 |
| Agar BSB | 4.25±0.03$^{a}$ | 6.27±0.12$^{b}$ | 1.33±0.10$^{b}$ | 48.8±0.44$^{a}$ | 130.6±15.7$^{a}$ | 3 | 2.04±0.39$^{b}$ | 4.22±1.44$^{a}$ | 0.41±0.12$^{a}$ | 19.9±3.48$^{c}$ | 2.17±0.71$^{a}$ | 20 |
| Liquid B | 4.04±0.06$^{d}$ | 7.1±1.08$^{b}$ | 1.19±0.14$^{b}$ | 36.3±4.78$^{c}$ | 355.9±273.4$^{a}$ | 3 | 1.13±1.10$^{bc}$ | 1.71±1.42$^{bc}$ | 0.29±0.14$^{b}$ | 20.8±8.70$^{bc}$ | 1.23±0.53$^{cd}$ | 18 |
| Liquid BS | 4.12±0.05$^{bcd}$ | 8.6±0.17$^{a}$ | 1.67±0.18$^{a}$ | 36.6±1.30$^{bc}$ | 291.9±94.8$^{a}$ | 3 | 1.75±0.78$^{bc}$ | 2.07±1.02$^{b}$ | 0.26±0.06$^{b}$ | 27.9±3.29$^{ab}$ | 0.77±0.24$^{e}$ | 22 |
| Liquid BSB | 4.18±0.06$^{abc}$ | 9.9±0.26$^{a}$ | 1.87±0.03$^{a}$ | 43.2±1.33$^{ab}$ | 240.0±21.5$^{a}$ | 3 | 1.86±0.51$^{bc}$ | 3.37±1.71$^{a}$ | 0.40±0.08$^{a}$ | 28.8±3.42$^{a}$ | 1.38±0.45$^{c}$ | 21 |
| Agar | 4.18±0.08$^{a}$ | 5.68±1.21$^{b}$ | 1.12 ± 0.18$^{b}$ | 45.8 ± 2.87$^{a}$ | 131.5±39.3$^{b}$ | 9 | 2.09±0.77$^{a}$ | 3.2±1.72$^{a}$ | 0.29±0.14$^{a}$ | 19.3±4.96$^{b}$ | 1.68±0.75$^{a}$ | 63 |
| Liquid | 4.11±0.07$^{a}$ | 8.53±1.34$^{a}$ | 1.58 ± 0.18$^{a}$ | 38.7 ± 4.25$^{b}$ | 271.5 ± 156.6$^{a}$ | 9 | 1.77±0.81$^{b}$ | 0.86±0.63$^{b}$ | 0.32±0.11$^{a}$ | 27.5±6.03$^{a}$ | 1.18±0.51$^{b}$ | 61 |
| B | 4.13±0.07$^{b}$ | 5.62±1.78$^{a}$ | 0.97 ± 0.25$^{b}$ | 40.2 ± 5.57$^{a}$ | 182.2 ± 213.5$^{a}$ | 6 | 1.10±0.91$^{b}$ | 0.92±1.15$^{c}$ | 0.24±0.13$^{c}$ | 18.9±7.61$^{b}$ | 1.09±0.56$^{b}$ | 39 |

| | | | | | | | | | | | | |
|---|---|---|---|---|---|---|---|---|---|---|---|---|
| BS | 4.11±0.07[b] | 7.62±1.12[a] | 1.48±0.25[a] | 40.5±4.47[a] | 165.2±110.3[a] | 6 | 1.90±0.47[a] | 2.99±1.2[b] | 0.28±0.08[b] | 24.4±6.40[a] | 1.11±0.61[b] | 44 |
| BSB | 4.21±0.07[a] | 8.08±2.0[a] | 1.60±0.30[a] | 46.0±3.21[a] | 181.0±58.5[a] | 6 | 2.18±0.81[a] | 3.76±1.58[a] | 0.41±0.10[a] | 24.5±5.11[a] | 1.74±0.71[a] | 41 |

8

**Supplementary Text S1**. Fermentative parameters.

The calculation of standard fermentation performance parameters enables to frame this results in a biotechnological context by evaluating process efficiency independently of the tested scale. The highest biomass concentration ($C_x$; **Supplementary Table S1**; **Supplementary Equation S1**) occurred mid-way in the lowest volume (11.20 $gL^{-1}$), while the lowest was recorded at the beginning of the largest culture volume (0.74 $gL^{-1}$). The highest biomass concentration relative to volume shifted from day 9 to day 14 as volume increased, following the maximum dry biomass of the mats (**Supplementary Table S1; Figure 2B**). The highest volumetric productivity ($Q_x$; **Supplementary Equation S2**) of biomass was also found in the smallest volume (0.06 $gL^{-1}h^{-1}$), peaking at day 7, indicating that within this volume, nutrients and time were used more efficiently. Maximum productivity relative to volume tended to match the time of local maximum biomass concentration, meaning that, resource efficiency was highest per volume when biomass peaked.

$$C_x = \frac{pellicle\ biomass - inoculum\ biomass\ (g)}{volume\ (L)}$$

**Supplementary Equation S1**. Formula for biomass concentration ($C_x$), accounting for accumulated biomass and culture volume.

**Supplementary Equation S2**. Formula for volumetric productivity ($Q_x$), accounting for biomasss concentration and fermentation time.

The highest rate of substrate consumption ($Q_s$; **Supplementary Table S1**; **Supplementary Equation S3**) always coincided with the beginning of the analysis in all volume conditions, ranging between 0.1 $gL^{-1}h^{-1}$ and 0.07 $gL^{-1}h^{-1}$. This indicates that cultures were more metabolically active before day 7 than during subsequent time points, irrespective of culture volume. Accordingly, once a continuous pellicle was established, the rate of nutrient consumption declined. These observations support the correspondence between microbial growth phases and the distinct biological phenomena observed under LSSF conditions. Radial expansion at the air–liquid interface was associated with elevated metabolic activity, consistent with exponential growth. Following complete pellicle formation, the stabilization of metabolic activity reflected the transition toward the stationary phase, characterized primarily by biomass maintenance rather than rapid accumulation. As demonstrated by the model of glucose consumption rate, maintenance of mycelium mats tended to have a similar consumption irrespective of biomass, volume and age. Therefore, the gradual decrease in $Q_s$ over time corresponds to the averaging of this maintenance consumption rate and the heavy metabolically active period of radial expansion. Remarkably, biomass peaks consistently occurred after glucose depletion, most notably in lower volumes. The storage of energetic molecular reserves (such as glycogen) or the assimilation of nutrients that were not yet converted into structural fungal tissue can explain this delay in biomass peak.

$$Q_s = \frac{Consumed\ glucose\ (gL^{-1})}{Fermentation\ time\ (h)}$$

**Supplementary Equation S3**. Formula for substrate consumption rate ($Q_s$), accounting for consumed glucose and fermentation time.

The maximum biomass yield ($Y_{x/s}$; **Supplementary Table S1**; **Supplementary Equation S4**) overlapped with the time of highest biomass nearly across all volumes, shifting from day 9 up to day 14 as volume increased. The specific yield ranged between 0.07 $gg^{-1}$ to 0.57 $gg^{-1}$, meaning that for each gram of glucose the fungus produced a maximum of 0.57 g of fresh fungal biomass. The highest biomass yields were observed in the smallest culture volumes, particularly around the biomass peak. In contrast, larger volumes consistently exhibited lower and steadily increasing yields throughout the entire incubation period. As only pellicle biomass was harvested, the initial growth phase in suspension was not included in yield calculations. Because smaller volumes reached surface colonization earlier, glucose consumption associated with suspended growth was minimized. In contrast, larger volumes stayed in the first phase of growth in suspension for longer time, leading to a larger portion of consumption that was not accounted for in biomass quantification. This likely explains the twofold higher biomass yields observed in the lower-volume treatments.

$$Y_{x/s} = \frac{C_x\ (gL^{-1})}{Consumed\ glucose\ (gL^{-1})} \times 100$$

**Supplementary Equation S4**. Formula for biomass yield coefficient ($Y_{x/s}$), accounting for biomass concentration and consumed glucose.
